# Hypoxia promotes osteogenesis via regulation of the mito-nuclear communication

**DOI:** 10.1101/2022.02.26.482117

**Authors:** Andromachi Pouikli, Monika Maleszewska, Swati Parekh, Chrysa Nikopoulou, Juan-Jose Bonfiglio, Constantine Mylonas, Tonantzi Sandoval, Anna-Lena Schumacher, Yvonne Hinze, Ivan Matic, Peter Tessarz

## Abstract

Bone marrow mesenchymal stem cells (MSCs) reside in a hypoxic niche that maintains their differentiation potential. Several studies have highlighted the critical role of hypoxia (low oxygen concentration) in the regulation of stem cell function, reporting differentiation defects following a switch to normoxia (high oxygen concentration). However, the molecular events triggering changes in stem cell fate decisions in response to high oxygen remain elusive. Here, we study the impact of normoxia in the mito-nuclear communication, with regards to stem cell differentiation. We show that normoxia-cultured MSCs undergo profound transcriptional alterations which cause irreversible osteogenesis defects. Mechanistically, high oxygen promotes chromatin compaction and histone hypo-acetylation, particularly on promoters and enhancers of osteogenic genes. Although normoxia induces rewiring of metabolism, resulting in high acetyl-CoA levels, histone hypo-acetylation occurs due to trapping of acetyl-CoA inside mitochondria, likely due to lower CiC activity. Strikingly, restoring the cytosolic acetyl-CoA pool via acetate supplementation remodels the chromatin landscape and rescues the osteogenic defects. Collectively, our results demonstrate that the metabolism-chromatin-osteogenesis axis is heavily perturbed in response to high oxygen and identify CiC as a novel, oxygen-sensitive regulator of MSC function.

## INTRODUCTION

Mesenchymal stem cells (MSCs) are somatic stem cells that reside in various embryonic and adult tissues and can differentiate into bone, fat, cartilage, tendon and other organ progenitor cells, contributing to the organisation and maintenance of tissue integrity (Gomez-Salazar et al., 2020). Given their role in maintaining tissue homeostasis, it is not surprising that MSCs have been extensively used in regenerative medicine, as a promising tool during stem cell therapies (Parekkadan and Milwid, 2010). However, their clinical use requires extended *in vitro* expansion, which alters their physiological properties. More precisely, the absence of niche molecular signals and the lack of interaction with neighbouring cells have been found to affect several parameters of the stem cell behaviour during *in vitro* culture of MSCs (Gomez-Salazar et al., 2020). Furthermore, oxygen concentration has a dramatic impact on stem cell biology; indeed, the MSC niche within the bone marrow cavity is characterised by low oxygen tension, ranging from 1.3% to 4.2% O2 (Spencer et al., 2014). However, MSCs used in research studies as well as in therapeutics are usually cultured under normoxic conditions, i.e. 21% O2, which significantly affect their activity (Parekkadan and Milwid, 2010). In fact, several reports have demonstrated that high oxygen drives stem cell fate decisions and impairs commitment of MSCs to specific lineages (Buravkova et al., 2014; Fehrer et al., 2007). However, we still lack a comprehensive understanding of the molecular, cell-intrinsic mechanisms linking oxygen concentration to MSC function.

Cells rewire their metabolism in response to oxygen availability and these metabolic changes influence stem cell differentiation capacity (Leijten et al., 2014); in turn, efficient differentiation of MSCs into specific cell types is directly associated with oxygen availability and requires profound metabolic rearrangements, such as down-regulation of glycolysis and induction of oxidative phosphorylation (OxPhos) (Shyh-Chang et al., 2013). Furthermore, metabolism is tightly linked to the epigenome, via central metabolites that serve as cofactors and substrates for epigenetic enzymes (Dai et al., 2020; Etchegaray and Mostoslavsky, 2016; Lu and Thompson, 2012). Therefore, oxygen-driven changes in cell metabolism could directly alter the epigenome of stem cells. Notably, the epigenetic landscape determines the transcriptional output, which regulates stem cell fate decisions and controls stem cell identity. Thus, characterising in depth the metabolism-chromatin-stemness axis could help develop clinical strategies to intervene to MSC fate decisions, aiming at enhancing their therapeutic potential. Here, we used bone-marrow-derived MSCs to investigate the role of oxygen in the regulation of stem cell fate decisions. We found that high oxygen leads to defective osteogenesis and impaired MSC function due to changes in the transcriptional profile. Mechanistically, high oxygen induces epigenetic alterations characterised by chromatin compaction and histone hypo-acetylation, particularly on promoters and enhancers of osteogenic genes. Despite the oxygen-driven metabolic rewiring which results in production of high acetyl-CoA levels, histone hypo-acetylation occurs due to trapping of acetyl-CoA inside the mitochondria of normoxia-cultured cells, indicating that the functionality of citrate carrier (CiC) is impaired at high oxygen. Remarkably, similar changes were observed in ageing MSCs, suggesting that high oxygen mimics the physiological process of ageing with regards to the role of the mito-nuclear communication in the regulation of stem cell differentiation (Pouikli et al., 2021).

## RESULTS

### Normoxia impairs osteogenesis via altering the MSC transcriptome

In order to investigate the role of oxygen concentration in the metabolism-chromatin-stem cell fate axis, we used freshly isolated bone marrow MSCs. We isolated and cultured MSCs based on a protocol that allowed purification of a highly homogeneous and tri-lineage potent population of MSCs which expressed increased levels of the Sca-1 and PDGFRa markers, maintained throughout the *in vitro* cell culture period (Pouikli et al., 2021). The sorted MSC population was then divided in two groups; one was transferred back to hypoxic conditions, whereas we shifted the other one to normoxia, as outlined in **Figure 1a**. In order to allow cells to adapt in the high oxygen environment, we performed all experiments seven days after transferring cells to normoxia, thus focusing on the long-term effects of normoxia in stem cell function. Culturing cells under normoxic conditions did not impact their adipogenic differentiation capacity (**Figure 1b-1c)**.

**Figure 1.**
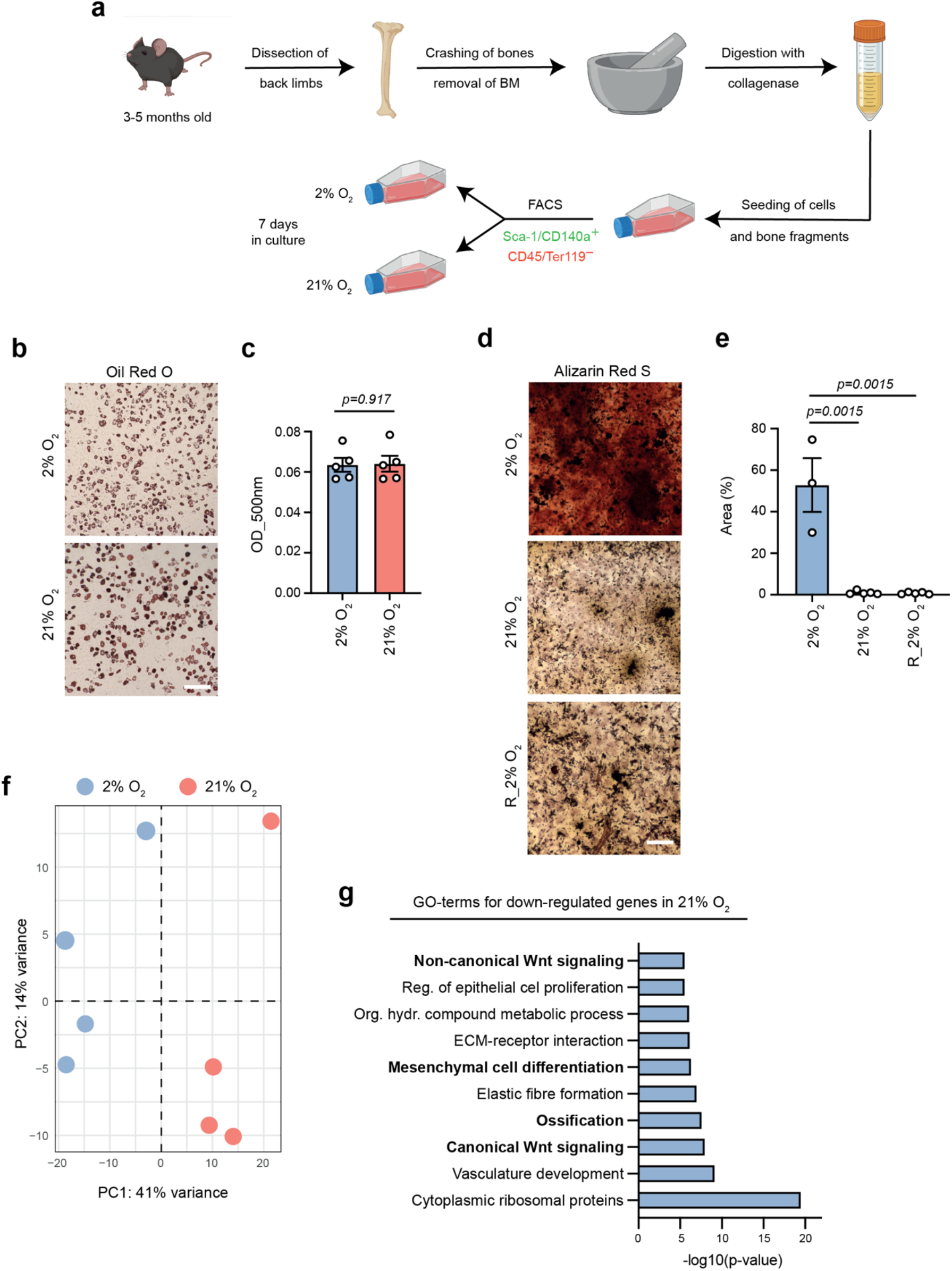
Normoxia changes the transcriptional output to suppress osteogenesis. (**a**) Schematic representation demonstrating the isolation protocol and the culture conditions of BM-MSCs isolated from the back limbs of young (∼3-5 months old) mice. After collection of the limbs, clean bones were cut into small pieces, which were then treated with collagenase for 1 hour at 37°C. Cells and bone fragments were seeded in flasks and incubated for 10 days under 2% O_2_. On day 10, cell sorting was performed using flow cytometry and selecting the CD45-/Ter-119-/Sca-1+/CD140a+ mesenchymal stem cell population. The isolated population was then split into two groups: one was transferred back to hypoxia, whereas the other one was shifted to normoxia. Cells were cultured under these conditions for 7 days. (**b-c**) Representative images (*b*) and quantification (*c*) of Oil Red O Staining of hypoxia- and normoxia-cultured cells, 9 days after induction of adipogenesis. n=5 biologically independent replicates. (**d-e**) Representative images (*d*) and quantification (*e*) of Alizarin Red S staining of hypoxic, normoxic and reversed hypoxic (R_2% O_2_) cells, 12 days after induction of osteogenesis. Cells were exposed to 21% O_2_ for 7 days and then moved back to 2% O_2_, where osteogenesis was induced after 4 days. (**f**) Principal component analysis (PCA) plot showing clustering of hypoxia- and normoxia-cultured cells after RNA-seq. n=4 biologically independent replicates. (**g**) GO enrichment analysis for down-regulated genes upon exposure to normoxia, as identified by RNA-seq. Scale bars, 500μm.

However, we found that normoxia-cultured cells exhibited profound defects in their osteogenic differentiation potential (**Figure 1d-1e)**, strongly indicating that normoxia promoted skewed adipogenic differentiation at the expense of osteogenesis. Furthermore, we demonstrated that normoxia impaired osteogenesis permanently, since shifting MSCs back to hypoxia, after culture under normoxic conditions for seven days, did not rescue the impaired osteogenic differentiation (**Figure 1d-1e)**. Interestingly, the transcriptomic profile of MSCs was also heavily influenced by the high oxygen levels, with normoxia-cultured cells clustering separately from those maintained permanently under hypoxic conditions (**Figure 1f**). Notably, GO term analysis revealed that the down-regulated genes in normoxia were involved in processes and signalling cascades associated with bone development and osteogenesis (**Figure 1g**). This suggests that high oxygen induced profound alterations in the transcriptional output of MSCs leading to reduced osteogenesis.

### High oxygen promotes global chromatin compaction and increased nucleosomal density on osteogenic gene promoters

Given that chromatin architecture is involved in the regulation of gene transcription, we next sought to investigate whether the normoxia-driven changes in the transcriptional profile were dictated by alterations in the chromatin landscape. Initially, to test whether exposure to high oxygen influenced chromatin accessibility in MSCs, we performed ATAC-seq on hypoxia- and normoxia-cultured cells (**Extended Figure 1a-1d)**. PCA analysis showed that cells were clearly distinct and separated based exclusively on the oxygen conditions where they were cultured (**Figure 2a)**. Furthermore, we identified, in total 81,393 accessible sites of which 23,571 changed significantly (FDR<0.01) in accessibility upon shift to atmospheric oxygen (**Figure 2b);** in particular 7857 sites became more open whereas 15714 sites became more compact.

**Figure 2.**
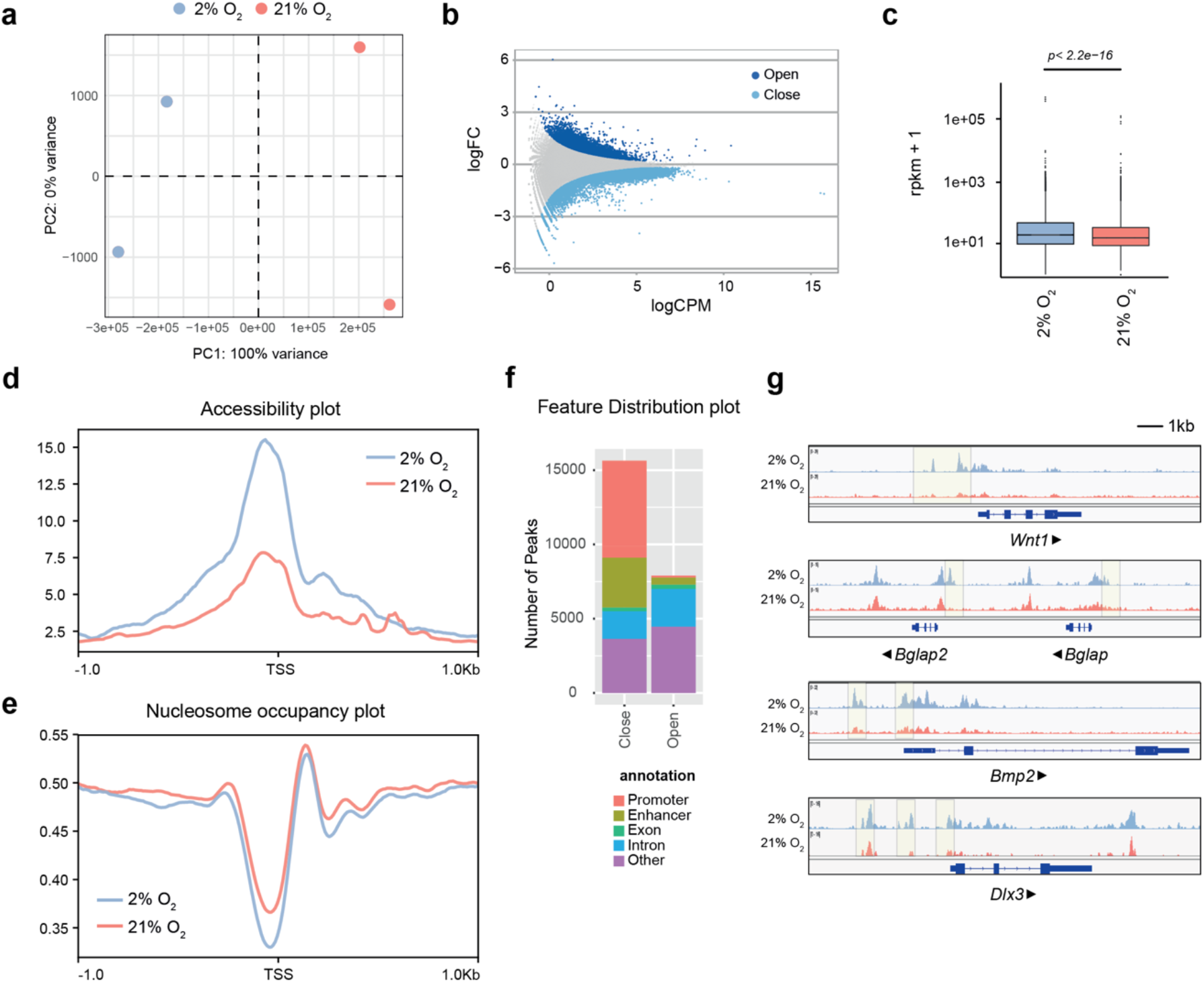
Normoxia leads to reduced chromatin accessibility. (**a**) Principal component analysis (PCA) plot showing clustering of hypoxia- and normoxia-cultured cells after ATAC-seq. (**b**) MA-plot showing opening and closing peaks upon shift to high oxygen, as determined by ATAC-seq. (**c**) Overall genome accessibility expressed as RPKM values measured by ATAC-seq. The y axis is scaled to log10 for visualization purposes. (**d**) Metaplot of ATAC-seq reads over the TSS of all protein-coding genes. (**e**) NucleoATAC metaplot to map position of all nucleosomes around the TSS of all protein-coding genes. (**f**) Feature distribution plot showing that most differentially accessible sites fall into promoters and enhancers. (**g**) Integrative genomics viewer (IGV) browser views showing ATAC-seq read density in hypoxia- and normoxia-cultured MSCs, near the *Wnt1, Bglap, Bmp2* and *Dlx3* gene promoters. Shaded regions demonstrate accessibility differences between hypoxia- and normoxia-cultured cells. Arrows indicate direction of transcription. n=2 biologically independent experiments.

Thus, we demonstrated that normoxia led to global chromatin compaction (**Figure 2c)**. We further analysed differences in the chromatin structure and plotted accessibility over the transcription start site (TSS), as a metaplot. We observed that gene promoters of cells exposed to high oxygen exhibited a strong decrease in chromatin accessibility (**Figure 2d)**. To validate this result, we employed NucleoATAC, an algorithm that allows the precise mapping of nucleosomes from ATAC-seq datasets (Schep et al., 2015). Plotting nucleosome occupancy over the TSS of all protein-coding genes confirmed that atmospheric oxygen resulted in higher nucleosome density around the promoter region, but more importantly, confirmed the reduced chromatin accessibility at the TSS (**Figure 2e)**. Feature distribution analysis, using promoter proximity criteria to call gene promoters and previously published enhancer regions in MSCs (Pouikli et al., 2021) to identify potential enhancers, confirmed that the vast majority of closing peaks in normoxia-cultured cells were found in gene regulatory elements i.e., promoters and enhancers (**Figure 2f)**. This strong decrease in chromatin accessibility at the TSS was also visible on a single gene level. For instance, promoters of genes important for osteogenesis, such as *Wnt1, Bglap, Bmp2* and *Dlx3*, showed decreased chromatin accessibility in normoxia-cultured cells (**Figure 2g**). Overall, osteogenic promoters were dramatically remodelled in response to high oxygen, strongly indicating that the transcriptional and functional osteogenesis defects were regulated on the chromatin level.

### Normoxia leads to reduced histone acetylation

To further investigate the molecular underpinnings for the normoxia-associated chromatin compaction, we compared histone acetylation levels in hypoxia- and normoxia-cultured cells. Histone acetylation neutralises the positive charge of the side-chain of histones and weakens the contact between histones and DNA (Tessarz and Kouzarides, 2014). Thus, it plays a fundamental role in the regulation of chromatin accessibility. Therefore, we investigated if alterations in the histone acetylation profile contributed to normoxia-induced changes in chromatin accessibility. We initially followed an unbiased approach and performed quantitative SILAC-MS, as outlined in **Figure 3a**, using heavy-labelled commercially available MSCs as SILAC spike-in standard. Although the coverage of the detected modifications was not extensive due to low cell numbers, our results clearly demonstrated that while histone methylation remained largely unaffected, normoxia-cultured cells displayed reduced histone acetylation (**Figure 3b**). In line with this, culture of MSCs under normoxic conditions led to lower histone H3 acetylation levels, as evidenced by immunofluorescence experiments (**Figure 3c-3d**).

**Figure 3.**
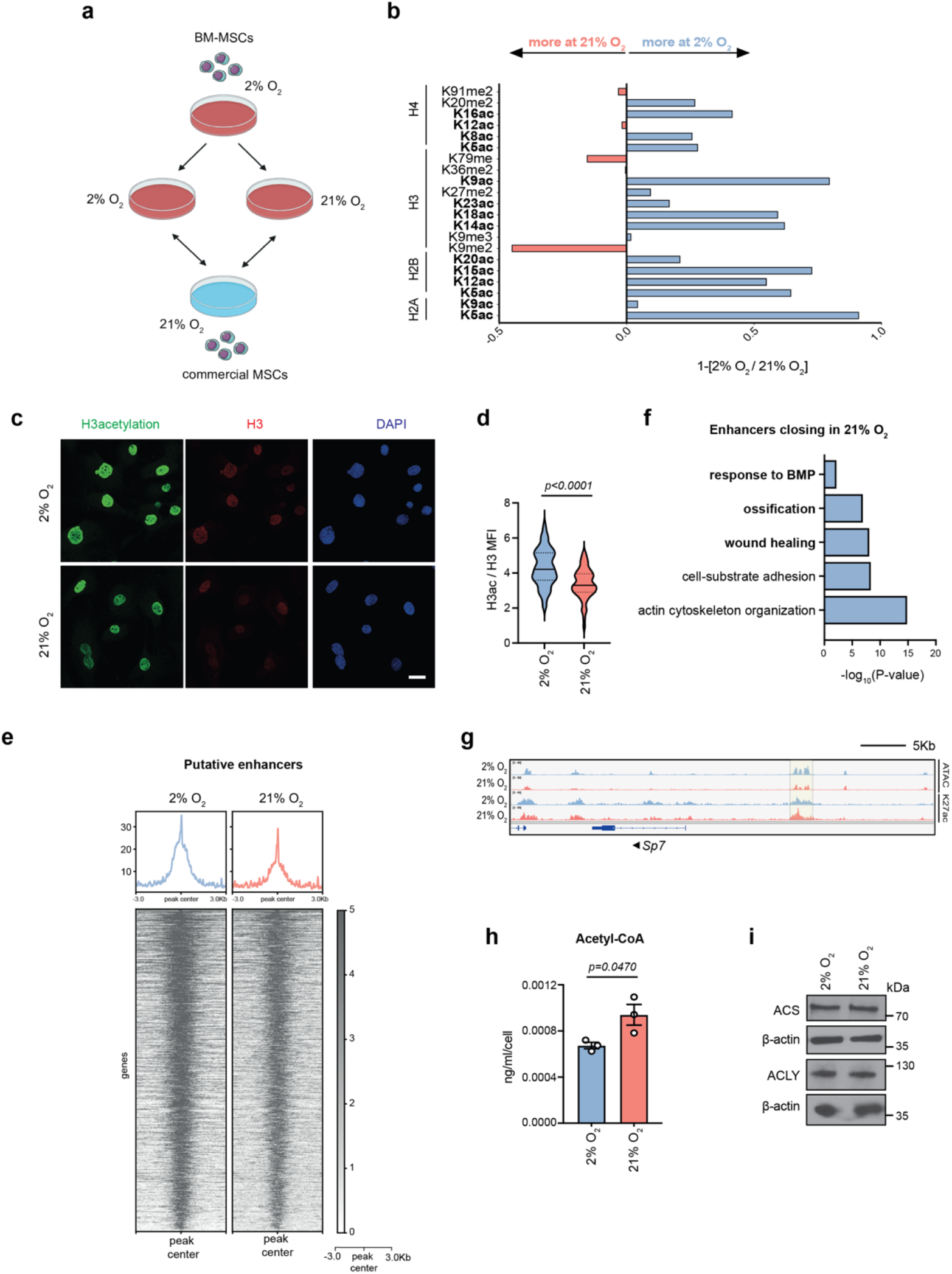
Normoxia results in histone hypo-acetylation on osteogenesis-involved enhancers. (**a-b**) Schematic representation demonstrating the protocol followed in the SILAC-MS experiment, where hypoxia- and normoxia-cultured cells were mixed with heavy-labelled commercial MSCs (*a*) and analysis of the levels of the detected histone modifications (*b*). n=2 biologically independent experiments. (**c-d**) Representative images and quantification of Mean Fluorescence Intensity (MFI) after immunostaining against H3ac and H3 of hypoxic and normoxic cells. MFI of histone H3 was used as internal control, for normalization. Nuclei were stained with DAPI. n=3 biologically independent experiments and results of a representative experiment are shown in *d*. Scale bar, 25μm. (**e-f**) Heatmaps showing chromatin accessibility at putative enhancers, in hypoxia- and normoxia-cultured cells. (**g**) Integrative genomics viewer (IGV) browser views showing H3K27ac read density in hypoxia- and normoxia-cultured MSCs, likely in the *Sp7* gene enhancer and promoter regions. Shaded region demonstrates differences in H3K27ac abundance between hypoxia- and normoxia-cultured cells. Arrow indicates direction of transcription. n=2 biologically independent experiments. (**h**) Liquid chromatography Mass spectrometry (LC-MS) analysis of acetyl-CoA in hypoxia- and normoxia-cultured cells. n=3 biologically independent experiments. (**i**) Representative immunoblots for ACS and ACLY in hypoxia- and normoxia-cultured cells. β-actin was used as loading control. n=3 biologically independent experiments.

Given the strong impact of high oxygen on enhancer accessibility **(Figure 2f)**, we focused on oxygen-induced changes in the functionality of potentially regulated genes. First, we generated ChIP-seq data for H3K27 acetylation (H3K27ac), a mark associated with active enhancers, using hypoxia- and normoxia-cultured cells. Integration of H3K27ac and ATAC-seq datasets confirmed the overall decrease in enhancer accessibility in normoxia-cultured MSCs (**Figure 3e**). Subsequently, we annotated genes on the basis of the ‘closest distance’ method (Moore et al., 2020). Focusing on the closing peaks in normoxia, it was apparent that decreased enhancer accessibility was associated with genes involved in MSC function, particularly in ossification and BMP signalling (**Figure 3f**). This indicated that loss of enhancer accessibility played a crucial role in the loss of the osteogenic potential in response to high oxygen levels. Example of an osteogenic gene which lost both chromatin accessibility and H3K27ac abundance on potential enhancer in normoxia is shown for *Sp7* (**Figure 3g**). Together our data indicated that high oxygen resulted in histone hypo-acetylation and chromatin condensation which led to lower osteogenic differentiation potential.

### Normoxia induces a metabolic rewiring which results in higher acetyl-CoA levels

Our findings prompted us to investigate how the histone acetylation profile was established in hypoxia- and normoxia-cultured MSCs and whether other cellular processes were involved in this phenotype. Thus, we next sought to characterise the metabolic profile of hypoxic and normoxic cells. Consistent to previous reports, we showed that hypoxia-cultured cells relied heavily on glycolysis for energy production (**Extended Figure 2a-2e**), whereas high oxygen resulted in stimulation of the mitochondrial activity and upregulation of OxPhos (**Extended Figure 2f-2h**). Unexpectedly, we found that this metabolic rewiring led to increased acetyl-CoA levels in normoxic MSCs (**Figure 3h**). Given that acetyl-CoA availability directly influences levels of histone acetylation, this result was in contradiction to the histone hypo-acetylation observed in MSCs from normoxia-cultured cells, but was similar to the phenotype observed in aged MSCs (Pouikli et al., 2021). Therefore, we set out to investigate further where the higher levels of acetyl-CoA originated from and what they were used for. One main metabolic pathway that heavily generates and consumes acetyl-CoA is lipid metabolism. To understand if the lipid content changed in response to high oxygen, we stained cells with Nile Red to visualise neutral lipids. We found that normoxic MSCs exhibited a much weaker lipid staining, indicating lower content of neutral lipids (**Extended Figure 3a-3b**). This observation was also confirmed with electron microscopy data that showed a strong decrease in the amount of lipid droplets (LDs) in normoxic MSCs (**Extended Figure 3a**). The lower LD content could be explained both by lower fatty acid biosynthesis and by increased lipid consumption through β-oxidation. Focusing on potential alterations in the lipid biogenesis pathway and comparing the levels of enzymes involved in *de novo* synthesis of fatty acids, we showed that the protein levels of the master regulator of lipogenesis SREBP1 and of the key-lipogenic enzymes FASN and ACC1 were strongly decreased in normoxic cells (**Extended Figure 3c-3d**). This result suggested that acetyl-CoA entrance into the lipid biogenesis pathway was inhibited in response to high oxygen. In sum, our findings indicated that normoxia influenced central metabolic pathways, leading to increased production of acetyl-CoA which accumulated in the cells.

### Acetyl-CoA is trapped with the mitochondria of normoxic cells

Despite the fact that normoxic cells contained higher acetyl-CoA levels, they exhibited global loss of histone acetylation (**Figure 2**). Therefore, we then investigated how this apparent contradiction is explained. One potential reason for the reduced lipid biogenesis and histone acetylation in normoxia could be a decrease in the cytosolic pool of acetyl-CoA.

However, levels of the enzymes ATP-citrate lyase (ACLY) and acetyl-CoA synthetase (ACS), which are both involved in the cytoplasmic/nuclear generation of acetyl-CoA (Mews et al., 2017; Wellen et al., 2009), were stable upon switch to normoxic conditions (**Figure 3i**). Hence, we speculated that in normoxia-cultured cells, acetyl-CoA might be trapped inside mitochondria leading to impaired cytosolic/nuclear acetyl-CoA pool which would in turn result in decreased lipogenesis and histone hypo-acetylation. To test this hypothesis, we stained hypoxia- and normoxia-cultured cells with a pan-acetylated antibody that recognizes all acetylated lysine residues, and we used TOMM20 as a counterstain for mitochondria. Of note, when acetyl-CoA accumulates in the mitochondria in excessive amounts, it promotes the non-enzymatic acetylation of mitochondrial proteins (James et al., 2017). Surprisingly, we found that upon culture of MSCs at high oxygen levels, mitochondria accumulated more acetylated proteins and the signal of the acetyl-Lysine antibody shifted from nuclear to mitochondrial (**Figure 4a-4b**). Recently, we demonstrated that ageing MSCs exhibit a similar phenotype and we identified that lower levels of citrate carrier (CiC) are responsible for the impaired export of acetyl-CoA from the mitochondria to the cytosol (Pouikli et al., 2021). Therefore, we next compared CiC levels in hypoxia- and normoxia-cultured cells. Expression levels of the CiC-encoding gene and CiC protein levels were comparable between the two culture conditions (**>Extended Figure 4f-4g**), suggesting that CiC levels were not influenced by oxygen concentration. However, we next perturbed CiC function, either by supplementing normoxic cells with the acetyl-CoA precursor sodium acetate, or by inhibiting CiC function in hypoxic cells using the BTA drug. We found that BTA treatment led to repression of lipogenesis (**Figure 4d-4e**) and accumulation of acetyl-CoA within the mitochondria of hypoxic cells (**Figure 4f-4h**), thus mimicking normoxia. By contrast, acetate supplementation of normoxic cells was sufficient to restore lipid biogenesis and nuclear localisation of acetyl-CoA (**Figure 4d-4h**), reverting the effects of high oxygen and mimicking hypoxic conditions. Since changes in CiC function directly influenced lipogenesis and histone acetylation, we concluded that the normoxia-cultured cells displayed impaired CiC function, which resulted in trapping of acetyl-CoA within the mitochondria.

**Figure 4.**
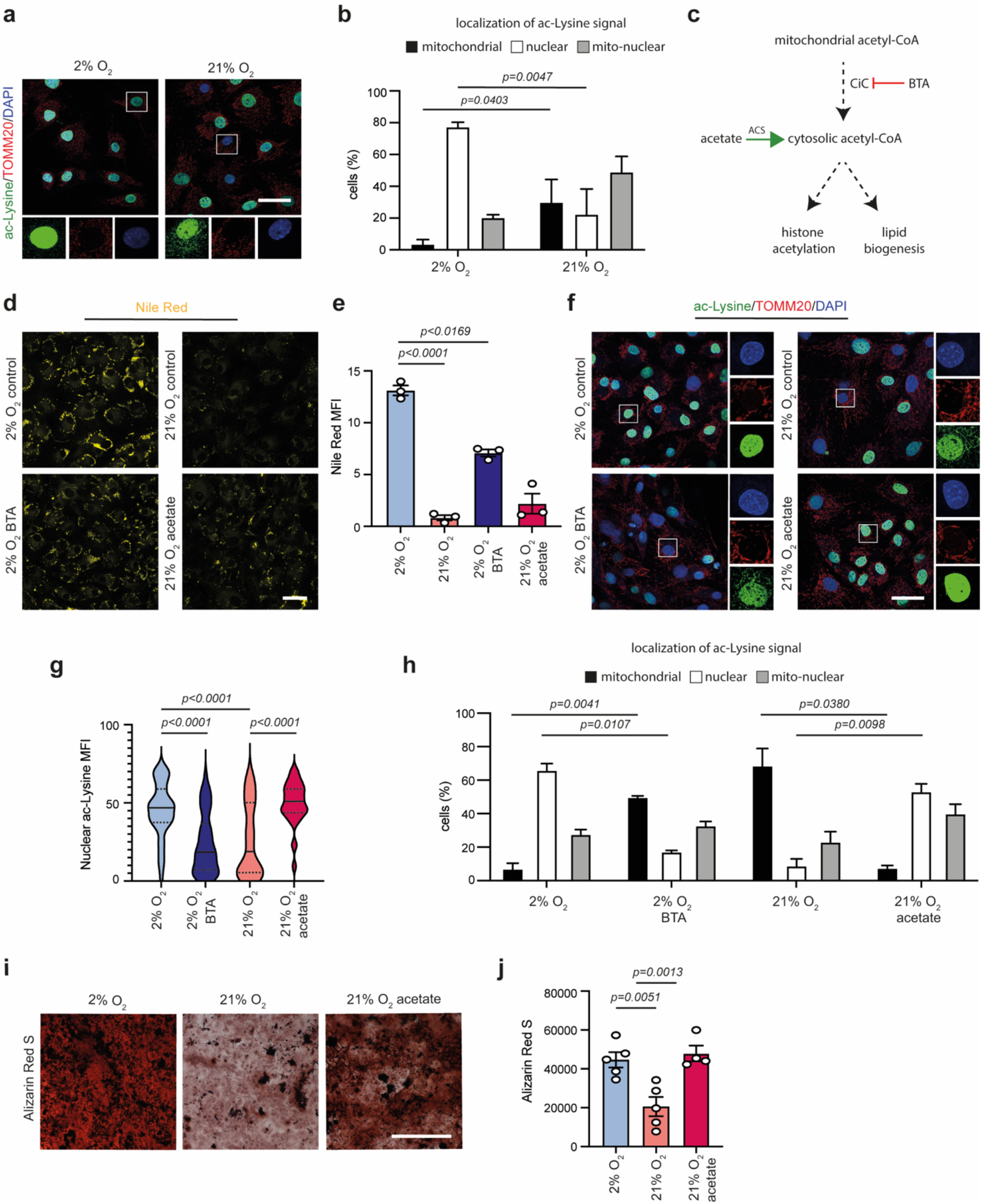
Trapping of acetyl-CoA inside mitochondria of normoxia-cultured BM-MSCs. (**a-b**) Representative images after immunostaining of hypoxia- and normoxia-cultured cells against acetyl-Lysine and TOMM20 (*a*), and assessment of acetyl-Lysine signal localization, after manual assignment into three categories: exclusively nuclear, exclusively mitochondrial and nuclear/mitochondrial (*b*). Nuclei were stained with DAPI. n=3 biologically independent experiments. Scale bar, 25μm. (**c**) Schematic graph illustrating the different pathways generating cytosolic acetyl-CoA and its potential routes. (**d-e**) Representative images (*d*) and quantification of MFI after Nile Red staining of hypoxic control and BTA-treated cells and of normoxic control and acetate-treated cells (*e*). Treatments were done for three days with 1mM BTA or 5mM acetate. Merged results of n=3 biologically independent experiments are shown. Scale bar, 25μm. (**f-h**) Representative images after immunostaining of hypoxic control and BTA-treated cells and normoxic control and acetate-treated cells against acetyl-Lysine and TOMM20 (*f*), quantification of nuclear acetyl-lysine MFI (*g*) and assessment of acetyl-Lysine signal localization, as described above (*h*). Nuclei were stained with DAPI. n=3 biologically independent experiments and results of a representative experiment are shown in *g*. Scale bar, 75μm. (**i-j**) Representative images and quantification of Alizarin Red S staining of control hypoxic and normoxic cells and of normoxic acetate-treated cells, 12 days after induction of osteogenesis. Merged results of n=5 biologically independent experiments are shown in *j*. Scale bar, 500μm.

Last, we sought to investigate the functional consequences of normoxia-induced loss of CiC activity. Since efficient export of acetyl-CoA from mitochondria to the cytosol is indispensable for lipogenesis, which is in turn required for membrane synthesis during proliferation and efficient commitment to osteocytes, we speculated that the decrease of CiC function was responsible for the observed changes in osteogenesis. To test this hypothesis, we pre-treated normoxic cells with acetate and we then induced osteogenic differentiation. Impressively, acetate pre-treatment of normoxic cells rescued their ability to differentiate into osteoblasts (**Figure 4i-4j**). Importantly, given that acetate was not present during the differentiation process and that the pre-treatment was sufficient to increase osteogenic differentiation potential, our data suggest that pre-treatment with acetate reset the chromatin landscape to make it more permissive for differentiation.

Together, our results show that the metabolism-chromatin-stemness axis is heavily affected by oxygen levels and identify CiC as a novel, oxygen-sensitive regulator of MSC function.

## DISCUSSION

In the present study we focused on the flux of acetyl-CoA from mitochondria to the nucleus, and we investigated how normoxia-associated changes in the flow of acetyl-CoA affect stem cell function. The data presented here suggest a model (**Figure 5**) whereby normoxia-driven changes in chromatin structure lead to transcriptional alterations that are responsible for the decreased osteogenic potential of MSCs cultured under high oxygen conditions. We show that, upon shifting cells from low to high oxygen, there is a switch in the subcellular localization of acetyl-CoA, which affects the epigenetic landscape. In particular, normoxic MSCs exhibit compartmentalised acetyl-CoA localization in mitochondria, as a result of impaired export to the cytosol, which might be explained by lower CiC activity. Strikingly, restoring histone acetylation, and thus chromatin plasticity, is sufficient to improve the impaired osteogenic capacity of normoxia-cultured MSCs, highlighting the fundamental role of CiC in the regulation of the metabolism–chromatin–osteogenesis axis in response to high oxygen.

**Figure 5:**
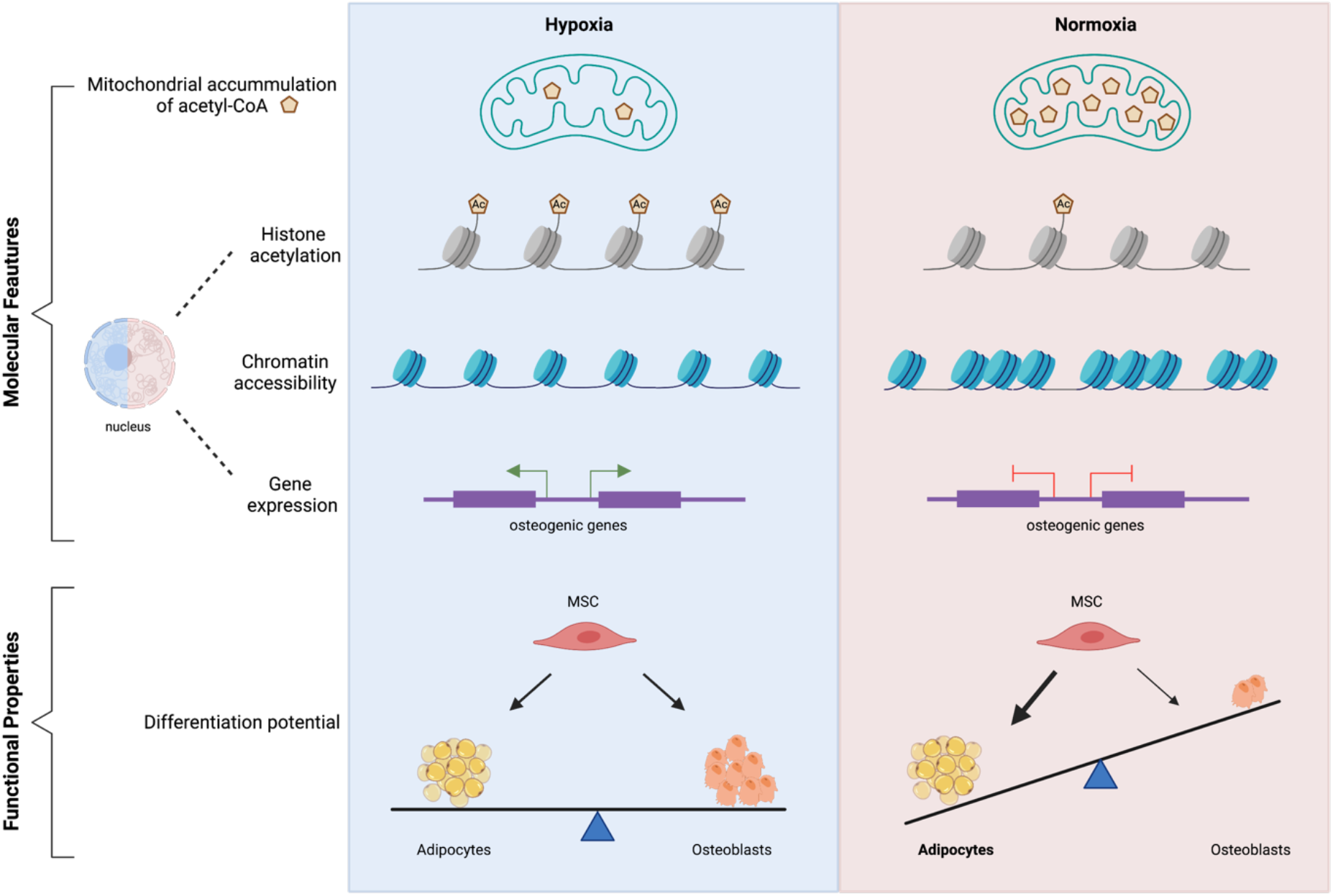
Model summarising the impact of shift to 21% oxygen. In the hypoxic environment acetyl-CoA is efficiently exported to the cytoplasm and can be used to acetylate histones, leading to a plastic chromatin state, which ensures balanced differentiation potential. Under atmospheric oxygen tension, acetyl-CoA remains trapped within mitochondria, which results in histone hypoacetylation, chromatin compaction and a shift in the differentiation potential favoring adipogenesis at the expense of osteogenesis.

The cellular environment plays a critical role in the regulation of stem cell fate decisions. Focusing on the role of oxygen tension on the metabolism-epigenome interplay with regards to stem cell differentiation, we report a negative impact of normoxia on the osteogenic potential of MSCs, which is in line to previous studies (Buravkova et al., 2014; Fehrer et al., 2007; Leijten et al., 2014; Pattappa et al., 2013). A recent report also highlighted that oxygen influences the metabolic state of MSCs and thus elicits changes in stem cell lineage commitment (Leijten et al., 2014). Consistent with this model, we identify that normoxia-cultured cells display reduced glycolysis (**Extended Figure 2a-2e**), which is usually associated with stemness and enhanced differentiation capacity. However, beyond the normoxia-driven metabolic alterations, we demonstrate that osteogenesis is also impaired due to epigenetic remodelling which leads to chromatin compaction and histone hypo-acetylation on promoters and enhancers of osteogenic genes (**Figures 2 and 3**). This, in turn, results in lower expression of these genes and irreversible defects in osteogenesis (**Figure 1d-1e)**. Whether changes in metabolism occur prior to the establishment of the epigenetic alterations and the extent to which each of these two pathways contributes to the reduced MSCs function is definitely worth further investigation. However, establishing a causal relationship might be extremely challenging due to the tight connection between chromatin and metabolism.

It has been shown that CiC function can be influenced by lysine acetylation. In particular, K97 acetylation increases CiC activity as it stabilises binding of its substrate citrate (Palmieri et al., 2015). However, several high-throughput studies have revealed many more potential acetylation (e.g. K255 (Weinert et al., 2013)) and phosphorylation (e.g. S156 (Tsai et al., 2015) sites within CiC, suggesting that CiC activity can be strongly modulated by post-translational modification (PTMs). Therefore, we speculate that although CiC levels are not impacted by high oxygen, CiC function might be compromised by specific normoxia-induced PTMs, leading to acetyl-CoA trapping within the mitochondria. Supporting the hypothesis of reduced CiC activity in response to high oxygen, modulation of CiC function, either by the inhibitor BTA or using acetate, directly influences histone acetylation and osteogenesis (**Figure 4f-4j**).

Strikingly, acetate supplementation of cells cultured under high oxygen is not only able to rescue histone acetylation levels, but also restores the osteogenic differentiation potential - similar to the situation in ageing MSCs (Pouikli et al., 2021). However, the impact of high oxygen on stem cell biology is much stronger than that of physiological ageing, as both chromatin accessibility and osteogenesis were more severely compromised under normoxic conditions. Notably, we performed these experiments after seven days of exposure to normoxia; while this is a time point when metabolic alterations have been already established, it might not be sufficient for the reset of other cellular processes. Importantly, MSCs isolated from mouse and human tissues, which have been adapted to atmospheric oxygen conditions, are proficient to differentiate into adipocytes, osteocytes and chondrocytes (Houlihan et al., 2012; Morikawa et al., 2009; Yang et al., 2018)). However, it is clear from the data reported here and previously published work that the differentiation potential is strongly enhanced under low oxygen conditions (Basciano et al., 2011; Georgi et al., 2015).

Of note, murine and human MSCs exhibit different extent of sensitivity to oxygen levels. Under low oxygen conditions, MEFs proliferate over an extended period of time. By contrast, at atmospheric oxygen levels, MEFs undergo senescence after a few doublings, whereas some cells escape this cell cycle arrest and start to proliferate as an immortal cell line (Parrinello et al., 2003). This spontaneous immortalisation correlates well with an increase in the accumulation of DNA damage. Importantly it is also known that mouse cells are more prone to acquire DNA lesions under normoxic conditions than cells derived from humans (Parrinello et al., 2003). We have not observed immortalisation of MSCs. Nevertheless, our findings indicate that an increase in oxygen concentration leads to phenotypes resembling physiological ageing. This raises the intriguing question of whether we investigate *de facto* aged cells in a lot of today’s models that we consider physiological and identifies oxygen concentration as a parameter that should be strongly considered when planning experiments using somatic stem cells as well as during stem cell therapies.

## MATERIALS AND METHODS

### Antibodies

For cell sorting, CD140a-APC (#17140181), Sca-1-FITC (#11598185), Terr-119-PE (#12592182) and the Fixable Viability Dye eFluor 450 (#65086314) were all purchased from eBioscience, whereas CD45-PE (#A16325) antibody was obtained from Life Technologies. For immunofluorescence and Western Blot experiments, antibodies against acetyl-Lysine (#9441S), Fatty Acid Synthase (#3189S), CBP (7389S), ACS (#3658T), anti-rabbit IgG HRP-linked (#7074S), anti-mouse IgG HRP-linked (#7076S) and histone H3 (#14269) were all purchased from Cell Signaling Technology. Antibodies against ACC1 (#21923-1-AP), ACLY (#15421-1-AP), KAT2A/GCN5 (14983-1-AP) and Citrate carrier (#15235-1-AP) were bought from ProteinTech. TOMM20 antibody (WH0009804M1) was obtained from Sigma-Aldrich. Histone H3 acetylation (#39139) and H3K27 acetylation (#39133) antibodies were all obtained from Active Motif. β-actin antibody (sc-47778) was bought from Santa Cruz Biotechnology. Anti-Mouse IgG Alexa fluor 488 (#A11001), anti-Rabbit IgG Alexa Fluor 488, anti-Rabbit IgG Alexa fluor 594 (#A-11012) were all bought from ThermoFisher Scientific.

## MOUSE MODEL AND EXPERIMENTAL DETAILS

### Mice

C57BL/6 N mice were bred and cared for in the mouse facility of the Max Planck Institute for Biology of Ageing. Mice were kept at a relative humidity of 50 ± 5%, a room temperature (RT) of 22 ± 2 °C and a light/dark cycle of 12 hours (6 a.m. to 6 p.m., with a 15-min twilight period). For our experiments we used exclusively wild-type male mice at 3–5 months.

### Endosteal-MSCs isolation

To isolate MSCs from their endosteal niche we followed a purification strategy based on a published protocol (Houlihan et al., 2012), and adapted the isolation strategy to acquire sufficient cells for our experiments. In brief, young C57BL6/N mice were sacrificed by cervical dislocation. Skin and muscles around the hind limbs were removed, the legs were cut above the pelvic joints and then placed in ice-cold PBS, on ice. Tibias were then separated from femurs by dislocating the joints and the clean bones were placed back in ice-cold PBS. All the following steps were performed inside the hypoxia hood. Bones were crushed and cut into tiny pieces and bone chips were incubated at 37°C for 75 minutes, in a-MEM medium containing 0.2% w/v collagenase (Sigma Aldrich), shaking at 200 rpm. To stop collagenase reaction, sample tubes with the bone chips were placed on ice and washed with culture medium (a-MEM supplemented with 10% FBS and 1% penicillin/streptomycin). We modified the published protocol by culturing the bone fragments together with the released cells; this allows the outgrowth of more MSCs from the bone *in vitro* before cell sorting by flow cytometry, increasing the cell yield we obtain. Bone chips were transferred into T25 flasks and were cultured under humidified conditions in hypoxia (2% O_2_, 5%CO_2_, 37°C). On day 3 of cell culture, the medium was changed and on day 5 both the cells and the bone chips were passaged, using Trypsin/EDTA solution (Life Technologies). On day 8 of the cell culture the bone chips were removed and on day 12 we performed cell sorting using flow cytometry.

### Cell sorting by Flow Cytometry

To obtain a purified MSC population we performed cell sorting by flow cytometry, using Fixable Viability Dye eFluor 450 (1:10000) and the following antibodies: Sca-1-FITC, CD140a-APC, CD45-PE and TER-119-PE, all in 1:1000 dilution. After harvesting, the cells were washed with PBS and resuspended in Hank’s Balanced Salt Solution (1x HBSS, 0.01M HEPES, 2% FBS, 1% penicillin/streptomycin, all Life Technologies). After incubation with the antibodies for 45 minutes, cells were washed twice with HBSS+ and were filtered through 35 μm nylon mesh into 5 ml sample tubes. Sorting was performed using BD FACSAria IIu and BD FACSAria IIIu instruments (BD Biosciences), under low pressure sorting conditions (100 μm nozzle and 20 psi sheath pressure). The CD45-/TERR-119-/CD140a+/Sca-1+ population was sorted into an eppendorf tube containing culture medium, at 4°C. Compensation was done using UltraComp compensation beads (#01222242, Invitrogen). Once sorted, MSCs were centrifuged at 300 g for 10 minutes at 4°C and resuspended in culture medium. Cells were cultured under humidified conditions (5% CO_2_, 37°C), in hypoxia (2% O_2_) and normoxia (21% O_2_), for seven days.

### qRT-PCR

Cells were lysed with QIAzol (QIAGEN) and total RNA was extracted using RNA extraction kit (Direct-zol RNA MiniPrep - Zymoresearch), following the manufacturer’s protocol. This was followed by cDNA synthesis using Maxima™ H Minus cDNA synthesis master mix (Thermo Scientific), according to the manufacturer’s instructions. Subsequent qRT-PCR was performed with 10 ng of cDNA, using SYBR-Green chemistry (Roche) on a Light Cycler 96 instrument (Roche). Data were analyzed and further processed in Microsoft Excel and Prism8 software. Fold change in gene expression over control samples was calculated using the ΔΔCq method, where β-actin Cq values were used as internal control. All reactions were run in three technical replicates and averaged. Experiments were performed three independent times and merged results are shown.

Oligos were designed using Primer3 and Blast platforms. The primers used for each gene are:

*Slc25a1 (encoding citrate carrier)*

5’ GGAGAGGACTATTGTGCGGTCT 3’

5’ CCCGTGGAAAAATCCTCGGTAC 3’

*Hk2*

5’ GTGGCTAGAGCTCGGGATC 3’

5’ TTTCCAGTCGCCCAACATCT 3’

Pgk1

5’ GCGCCACCTTCTACTCCTCC 3’

5’ CTTCCATTTGTCACGTCCTG 3’

*Pfk1*

5’ ACCGAATCCTGAGTAGCAAG 3’

5’ GTAGCCTCACAGACTGGTTC 3’

*Pgam1*

5’ ATCAGCAAGGATCGCAGGTA 3’

5’ TTCATTCCAGAAGGGCAGTG 3’

*β-actin*

5’ CTGCGCTGGTCGTCG 3’

5’ CACGATGGAGGGGAATACAG 3’

### Mitotracker staining

Cells were washed with PBS, harvested with Trypsin-EDTA (Life Technologies) and resuspended in the pre-warmed (37°C) staining solution containing the MitoTracker Deep Red FM probe (ThermoFischer Scientific #M22426) in a final dilution 1:15000 in assay medium (a-MEM without FBS and phenol red). Cells were incubated with the staining solution for 30 minutes at 37°C. They were then washed twice with PBS, centrifuged at 500 g for 5 minutes. and resuspended in assay medium. After addition of DAPI (Invitrogen) for dead-cell exclusion right before measurement, cells were analysed by flow cytometry (BD FACSCANTO II cytometer, BD Biosciences). Data was collected using FACS-Diva software and analysed using FlowJo software.

### Oxygen Consumption Rate

The SeaHorse XF96 extracellular Flux Analyzer (Agilent Technologies) was used to determine Oxygen Consumption Rate (OCR) in normoxia- and hypoxia-cultured MSCs. 2×10^4^ cells were seeded in 96-well SeaHorse plates, after coating them with 10% Gelatin - 90% Poly-L-Lysine solution for 1 hour. Cells were incubated overnight with culture medium in a humidified incubator. On the day of the experiment, cells were washed twice with assay medium (XF-DMEM, 10 mM Glucose, 1 mM Pyruvate, 2 mM L-Glutamine) and incubated for 1 hour, prior to loading into the XF Analyzer, in a non-CO_2_ - containing incubator. Following measurements of resting respiration, cells were injected subsequently with 20 μM oligomycin, 5 μM FCCP and 5 μM Rotenone/Antimycin (all drugs were from Agilent Technologies). Each measurement was taken over a 2-min interval followed by 2-min of mixing and 2-min of incubation. Three measurements were taken for the resting OCR: after oligomycin treatment, after FCCP and after Rotenone/Antimycin A treatment. To determine extracellular acidification rate (ECAR), 3×10^4^ cells were seeded in 96-well SeaHorse plates, as described above. On the day of the experiment, cells were washed twice with the assay medium (XF-DMEM, 2 mM L-Glutamine) and incubated for 1 hour in a non-CO_2_ - containing incubator, before loading into the XF Analyzer. Glycolytic activity was measured after injection with 10 μM Glucose, 1 μM Oligomycin and 50 μM 2-DG. Each measurement was taken over a 2-minute-interval followed by 2-minute-mixing and 2-minute-incubation. In both assays, values were normalised to protein concentration using Bradford kit and were plotted using the Wave 2.4 and the Prism8 software programmes.

### Differentiation assays

#### i) Differentiation to adipocytes

For adipogenesis experiments, 2×10^3^ cells were seeded in 96-well plates. Adipogenic differentiation was induced once cells reached confluency, by culturing them in culture medium or differentiation medium (culture medium supplemented with 1 μM dexamethasone, 1 μM IBMX, 10 μg/ml insulin, 100 μM indomethacin; all bought from Sigma Aldrich), for 8-10 days. Adipocytes were detected by Oil-Red-O (Sigma Aldrich) staining. Cells were washed with PBS and fixed in 3.7% formaldehyde (Roth) for 30 minutes, at RT. After fixation, cells were washed twice with ddH_2_O and once with 60% isopropanol (Roth), with every washing step lasting 5 minutes. Cells were then stained with Oil-Red-O staining solution for 15 minutes at RT. After staining, cells were washed four times with ddH_2_O and images were acquired with bright-field microscope, using the 20x objective.

#### ii) Differentiation to osteoblasts

For osteogenesis experiments, 2×10^3^ cells were seeded in 96-well plates. Osteogenesis was induced once cells reached confluency, by culturing them in culture medium or differentiation medium (culture medium supplemented with 100 nM dexamethasone, 10 mM beta-glycerophosphate, 100 μM ascorbic acid; all bought from Sigma Aldrich), for 11 days. For acetate treatment of normoxia cells, 5 mM of sodium acetate was added in the media three days prior to the induction of differentiation and was then removed. To observe whether osteogenic defects under normoxic conditions are reversible, cells were cultured in normoxia for 7 days, were then shifted to hypoxia and osteogenesis was induced after 4 days.

Osteoblasts were detected by Alizarin Red S (Sigma Aldrich) staining, performed 12 days after induction of osteogenesis. Cells were washed once with PBS and fixed in 3.7% formaldehyde for at least 30 minutes, at RT. Fixation was followed by washing of cells with ddH_2_O. Cells were then incubated with the Alizarin Red S staining solution (2% w/v Alizarin Red S in ddH_2_O) for 45 minutes, protected from light. After staining, cells were washed with ddH_2_O and images were acquired with bright-field microscope, using the 20x objective.

### Immunofluorescence experiments

For immunofluorescence experiments, 2×10^3^ cells were seeded in 96-well plates with glass bottom (Greiner) and treated as indicated in each experiment. After treatments, cells were fixed for 15 minutes at 37°C with 3.7% v/v formaldehyde in culture medium. Samples were washed with PBS twice, permeabilized with 0.1% TritonX-100 (Roth) in PBS for 10-15 minutes and blocked with 5% BSA in PBS (Roth) for 10 minutes. Samples were then incubated with the indicated primary antibodies diluted 1:100-1:300 in 3% BSA-PBS, overnight at 4°C. Following this, they were washed 3 times with PBS, with each washing step lasting 10 minutes. Samples were then incubated with the appropriate secondary fluorescent antibodies diluted 1:500 in 5% BSA-PBS for 45 minutes, protected from light. After 3 washing steps of 10 minutes each, cells were mounted using Roti-Mount FluorCare mounting medium (HP20.1, ROTH), containing DAPI. Images were acquired using 40x and 63x objective lens on SP8-X and SP8-DLS Leica confocal microscopes.

### Nile Red staining

1 mM of the Nile Red dye was added to the cells, diluted in HBSS+ buffer. Cells were incubated with the dye for 15 minutes at 37°C. They were then washed three times with HBSS+ buffer and live cell imaging was performed using the SP8-DLS Leica confocal microscope, with a 40x objective lens.

### Electron microscopy

Samples were fixed in fixation buffer (2% glutaraldehyde, 2.5% sucrose, 3 mM CaCl_2_, 100 mM HEPES pH 7.4), for 30 minutes at RT and 30 minutes at 4 °C. They were then washed with 0.1 M sodium cacodylate buffer (1% osmium, 1.25% sucrose, 10 mg/ml K_3_[Fe (CN)_6_] in 0.1 M sodium cacodylate), incubated for 1 hour on ice OsO4, and washed again with 0.1 M sodium cacodylate buffer. After ethanol washes, samples were incubated with EPON and embedded. Sections (70 nm) were cut using Leica Ultracut and put on negatively stained grids with 1.5 % uranylacetate in water, at 37 °C in the dark. Images were acquired using a JEM 2100 plus microscope (JEOL).

### Western Blot experiments

For all western blot experiments cells were harvested and lysed with RIPA buffer (150 mM NaCl, 1% TritonX-100, 0.5% sodium deoxycholate, 0.1% SDS, 50 mM Tris pH 8.0) supplemented with 5 mM sodium butyrate and 1x Protease Inhibitor Cocktail (Thermo Scientific). For efficient lysis, cells were incubated with RIPA buffer at 4°C, for 30 minutes, rotating, and then centrifuged for 10 minutes at 6500g. Protein concentration was determined using BCA protein assay kit (SERVA). 20-50 μg of total protein were loaded into each well and SDS-PAGE electrophoresis was performed at 160 V for ∼45 minutes. This was followed by transfer to nitrocellulose membrane, using the Trans-Blot Turbo blotting apparatus and reagents, all provided by Bio-Rad. Protein transfer was confirmed by Ponceau S staining (Sigma Aldrich) for 1-2 minutes. The membranes were then blocked using 5% non-fat dry milk in Tris-buffered saline-0.1% Tween20 (TBS-T) for 1 hour at RT. Membranes were incubated with the indicated primary antibodies, diluted in 3% Milk in TBS-T, at 4°C overnight, washed three times with TBS-T for 5 minutes each washing step and incubated with the appropriate horseradish peroxidase (HRP)-conjugated secondary antibodies diluted 1:10000 in 5% BSA in TBS-T, for 1 hour at RT. After three 10-minute washing steps in TBS-T, the desired proteins were visualised by providing fresh HRP-substrate solution (Luminol Enhancer Solution/Peroxide Solution - Promega) and exposure of membranes for specific time periods to photographic film, using the Curix60 instrument (Agfa).

### Measurement of glucose, lactate and pH

Measurement of glucose, lactate and pH in the media of MSCs was performed using the Vi-CELL MetaFLEX instrument (Beckman Coulter), according to manufacturer’s instructions. All measurements were done in triplicates and averaged.

### Targeted LC-MS analysis of acetyl-CoA

Metabolite extraction from each sample was performed using a mixture of 40:40:20 (v:v:v) of prechilled (−20 °C) acetonitrile:methanol:water (OptimaTM LC/MS grade, Thermo Fisher Scientific) and protein concentration was used for normalization (BCA Protein Assay Kit, Thermo Fisher Scientific). For analysis of acetyl-CoA, the extracted metabolites were resuspended in UPLC-grade acetonitrile:water (80:20 (v:v), OptimaTM LC-MS-grade, Thermo Fisher Scientific). The samples were analyzed on an Acquity iClass UPLC (Waters), using a SeQuant ZIC-HILIC 5 µm polymer 100 ×2.1 mm column (Merck) connected to a Xevo TQ-S (Waters) triple quadrupole mass spectrometer. The resuspended metabolite sample extract was injected onto the column and separated using a flow rate of 500 µl/min of buffer A (10 mM ammonium acetate, 0.1% acetic acid) and buffer B (acetonitrile) using the following gradient: 0–0.5 min 20% A; 0.5–1.4 min 20–35% A; 1.4–2.5 min 35–65% A. After 2.5 minutes the system was set back to 20% A and re-equilibrated for 2.5 minutes.

The eluted metabolites were detected in positive ion mode using ESI MRM (multireaction monitoring) applying the following settings: capillary voltage 1.5 kV, desolvation temperature 550 °C, desolvation gas flow rate 800 L/h, collision cell gas flow 0.15 ml/min. The following MRM transitions were used for relative compound quantification of acetyl-CoA m/z precursor mass (M+H+) 810, fragment mass (M+H+) m/z 303 using a cone voltage of 98 V and a collision energy of 28 V. For each compound, two further fragments were monitored as qualitative controls for compound identity. Data analysis and peak integration were performed using the TargetLynx Software (Waters).

### SILAC-MS analysis

#### SILAC Labelling

Commercially available murine MSCs (Cyagen, MUBMX-01001) were used as a standard for both, hypoxia and normoxia cultures. To label commercial MSCs, cells were grown in DMEM HG (w/o Lys and Arg, Thermo Scientific, # 88420), supplemented with 10% dialyzed FBS (Thermo Scientific, # 26400044) and heavy lysine (146mg/l; Sigma, # 608041) and arginine (28mg/l; Sigma # 608033) for a total of 6 passages. Total incorporation of heavy amino acids was determined by mass spectrometry to be >97%. Freshly isolated BM-MSCs were grown in the same medium, but with light lysine/arginine (Sigma, # L8662 and # A8094) for 7 days past FACS sorting at either 2% or 21% oxygen. Cells were subsequently mixed 1:1 and histones were isolated as follows.

#### Enrichment of histones

10^7^ cells of each replicate were harvested and spun down. Pellets were resuspended in 0.1 M H_2_SO_4_ and nutated for 2 hours at 4ºC. Solution was homogenized and subsequently centrifuged for 20 minutes at 3500 rpm, at 4ºC. Supernatant was neutralized with 1 M Tris-Cl pH 8.0 and the following reagents were added: 0.5 M NaCl, 2 mM EDTA, 0.25 mM PMSF, 1 mM DTT. Solution was added to a 2 ml SP-sepharose, washed with wash buffer (50 mM Tris pH 8, 0.6 M NaCl, 2 mM EDTA, 1 mM DTT, 0.25 mM PMSF) and eluted in elution buffer (50 mM Tris pH 8, 0.2 M NaCl, 2 mM EDTA, 1 mM DTT, 0.25 mM PMSF). Proteins were precipitated with 4% perchloric acid overnight at 4ºC and then spun at 14000 rpm, for 45 minutes at 4°C. Pellets were washed two times with 4% perchloric acid, two times with 0.2% HCl in acetone and two times with acetone. Pellets were dried and resuspended in ddH2O with 0.25 M PMSF.

#### Protein digestion

Proteins were digested using partial FASP as previously described (Leidecker et al., 2016). Briefly, proteins were resuspended in 8 M urea in 0.1 M Tris-HCl pH 8.0, 10 mM tris(2-carboxyethyl) phosphine (TCEP), 20 mM chloroacetamide and transferred to 10 kDa cut-off Vivacon® 500 flat filters. Samples were centrifuged at 14,000 g at 20ºC for 20 minutes, followed by three washes with 50 mM ammonium bicarbonate (ABC). For partial FASP digestion, 1:2000 trypsin Gold to protein ratio was used for 20 minutes at 20ºC. The digestion was stopped by the addition of formic acid to lower the pH below 3. Peptides were collected by centrifugation at 14,000 g at 4ºC for 10 minutes. Next, 50 mM ABC were added to the filter and peptides were collected by centrifugation at 14,000 g at 4ºC. This elution step was repeated. The retentate containing undigested proteins was further digested in 50 mM ABC overnight at 37ºC, with 1:50 trypsin to protein ratio. Peptides were collected by centrifugation at 14,000 g at 4ºC for 10 minutes, followed by a further elution with 50 mM ABC. Peptides were then desalted on either C18 cartridges (3M Empore) or using in-house manufactured StageTips (Rappsilber et al., 2003), depending on the peptide amounts. Eluted peptides were dried down in Speedvac concentrator and resuspended in 0.1% FA prior to LC-MS/MS analysis.

#### LC-MS/MS analysis

Liquid chromatography for all LC-MS/MS runs was performed on an EASY-nLC 1000 Liquid Chromatography system (Thermo Scientific) coupled to Q Exactive Plus mass spectrometer (Thermo Scientific) via modified NanoFlex sources (Thermo Scientific). Peptides were loaded onto 250-mm x 75-μm PicoFrit (C18 2 μm medium) analytical columns (New Objective) at a maximum pressure of 800 bar. Solutions A and B for the UPLCs were 0.1% formic acid in water and acetonitrile, respectively. Samples were loaded in 0.1% formic acid in water to maximize retention of highly hydrophilic peptides. Gradients varied slightly in length (90 to 150 min) and mixture, and may be extracted from the respective raw files. In general, they incorporated a linear gradient from very low or zero %B to 20 or 30% for 65-100 minutes, followed by a steeper phase and a wash. This length of gradient was maintained despite the relative simplicity of the protein mixture to improve the resolution and identification of as many modified peptide forms as possible, including those of low abundance. Full scan MS spectra were acquired from over an m/z range 300–1800 at 70 000 resolution, AGC targets were set to 3,000,000 ions, maximum injection time was 100 ms. MS2 acquisition varied slightly in resolution, AGC target and maximum injection time, and may be extracted from the respective raw files. In general, up to 5 data-dependent HCD fragmentation MS2 spectra were acquired at a resolution up to 70,000. AGC target for MS2 was set up to 1,000,000 ions. To reach this target, long MS2 injection times were allowed (up to 500 ms). Unassigned, singly-charged or >+8-charged ions were rejected and the dynamic exclusion option was enabled (duration: up to 40 s).

### Data analysis

Raw files were analysed with MaxQuant proteomics suite of algorithms (version 1.5.3.17) (Cox and Mann, 2008), integrated with the search engine Andromeda (Cox et al., 2011). The data were searched against a mice proteome database (downloaded 09.10.2015 from UniProt) with the following parameters: the maximum allowed mass deviation was set to 4.5 ppm for precursor ions and 20 ppm for fragment ions; the minimum peptide length was set to 6 amino acids; the maximum number of missed cleavages was set to 5 with the maximum charge state 6; multiplicity was set to 2 with Lys8/Arg10 as the Heavy Label and max. Labelled AAs were set to 7. Variable modifications included acetylation (Protein N-term and K), Methylation (KR), di-Methylation (KR), tri-Methylation (K) and Phosphorylation (STY). FTMS top peaks per 100 Da were set to 20.

### RNA-seq

Total RNA was isolated using the RNA extraction kit (Direct-zol RNA MiniPrep - Zymoresearch), following the manufacturer’s protocol. Once RNA quality and integrity were verified, RNA was submitted to library production at the Genomic Core Facility of the Max Planck Institute for Plant Breeding, Cologne using the NEBNext Ultra II RNA Library Prep Kit. Libraries were sequenced as single-end 150 bp reads on Illumina HiSeq 4000. The sequenced reads of RNA-seq dataset were processed using zUMIs (version 2.2.1) (Parekh et al., 2018) with STAR (version 2.6.1a) (Dobin et al., 2013), samtools (version 1.9) (Li et al., 2009) and featureCounts from Rsubread (version 1.32.4) (Liao et al., 2014). The reads were mapped to the mouse genome (mm10) with the ensembl annotation verion GRCm38.91. The generated count matrix was further analysed using R (version 3.5.1). First, genes were filtered using “filterByExpr” function of edgeR (Robinson et al., 2010), with the minimun count 5. The differential gene expression analysis between normoxic and hypoxic cells was carried out using limma-trend (Ritchie et al., 2015) approach at the adjusted p-value of 0.05. Obtained sets of genes were further analysed through gene ontology (GO) enrichment analysis.

### ATAC-seq

ATAC-seq was performed on 5×10^4^ cells per sample, as described previously (Buenrostro et al., 2013). DNA concentration of the libraries was measured using Qubit, and library quality was assessed by running samples on the TapeStation. Libraries were sequenced on a Illumina HiSeq 2500. The fastq files of sequenced reads were mapped to the mouse genome (mm10) using local alignment with bowtie2 (Langmead and Salzberg, 2012) with parameters -x mm10 and -X 2000. The resulting BAM files were sorted, indexed using samtools (version 1.3.1) and duplicates were removed using MarkDuplicates of Picard Tools. The peaks were called using chromstaR (Taudt et al., 2016) R package in differential mode between normoxic and hypoxic cells with the binsize of 300, stepsize of 100 and 15 as minimum mapping quality threshold. The per sample peak RPKM table was pulled out from the chromstaR model and differential accessibility analysis between normoxic and hypoxic cells was performed using edgeR (Robinson et al., 2010). The normalising factors were calculated using “RLE” method within “calcNormFactors”, tagwise dispersion trend was estimated using the default parameters in “estimateDisp” function and a generalised linear model was then fit on the data using “glmQLFit” function in robust mode. The peaks called by chromstaR were then used in nucleoATAC (Schep et al., 2015) for nucleosome positioning. For visualisation purposes, the replicates were merged using “samtools merge” and the bigwig files were generated using “bamCoverage --normalizeUsing RPGC”.

### ChIP-seq

ChIP-seq was performed as described previously (Tessarz et al., 2014), using 10^5^ MSCs per IP. The fastq reads were mapped to mm10 genome using bowtie2 (Langmead and Salzberg, 2012) and duplicates were then removed using MarkDuplicates program of Picard Tools. For mapping spike-in fragments to yeast, the --no-overlap --no-dovetail options were set and mapped to a repeat-masked version of the yeast genome (R64) to avoid cross-mapping of the mouse genome to that of the yeast genome. The peaks were then called using chromstaR (Taudt et al., 2016) package in differential mode between normoxic and hypoxic cells for H3K27ac with the binsize of 1000, stepsize of 500 and 15 as minimum mapping quality threshold. The differential analysis between normoxic and hypoxic cells for H3K27ac mark was performed using edgeR (Langmead and Salzberg, 2012) as described under the ATAC-seq analysis section. For visualisation purposes, the replicates were merged using “samtools merge” and the bigwig files were generated using “bamCoverage --normalizeUsing RPGC”.

### Image acquisition and processing

Bright-field images were acquired using the Evos FL Auto 2 microscope. Quantification of the Alizarin Red S staining was done in ImageJ software; measurement of intensity was done after making RGB stacks for each image and applying the same thresholds to all samples from each experiment. Immunofluorescent images were acquired using confocal SP8-X and SP8-DLS microscopes (Leica). All immunofluorescent images were processed identically in ImageJ; in particular, images are shown after background subtraction (rolling ball radius:50) and noise despeckle.

### Quantification and statistical analysis

Except for epigenome analysis, all other graphs were generated in GraphPad Prism8. For all bar graphs, results are shown as mean ± s.e.m. For quantification of images where more than ten cells were taken into account, distribution of data points is shown as violin plots, where the mean is indicated by a solid line and quartiles are indicated with dotted lines. *P*-values for box plots of sequencing data were determined using two-sided Wilcoxon test. Statistical significance was determined using two-sided unpaired *t*-test, unless otherwise specified, and the exact *P*-values are indicated on each plot. We performed GO enrichment analysis using Metascape, which calculates *P*-values on the basis of accumulative hypergeometric distribution.

## DATA AVAILABILITY

The mass spectrometry proteomics data have been deposited to the ProteomeXchange Consortium via the PRIDE (Perez-Riverol et al., 2019) partner repository with the dataset identifier XXXX. RNA- and ATAC-seq data is available at GEO, accession numbers: GSE 143580 and GSEYYYYY. Source data are provided with this paper.

## ACKNOWLEDGMENTS

The authors would like to thank members of the Tessarz lab for fruitful discussion on the impact of oxygen on cell physiology. Flow cytometry and fluorescent microscopy were performed at the FACS & Imaging Facility of the MPI for Biology of Ageing, Cologne. EM was performed at the Imaging facility of CECAD, University of Cologne. Sequencing was carried out at the Sequencing Core Facility of the MPI for Molecular Genetics, Berlin. This work was funded by the Max Planck Society (to P.T.), an Onassis foundation graduate fellowship (to A.P.) and an Alexander von Humbodlt postdoctoral fellowship (to M.M.). Models were drawn using Biorender.

## AUTHOR CONTRIBUTIONS

Conceptualization: A.P., M.M., P.T.; Experimentation: A.P., M.M., C.N., J.-J.B., Y.H.; Formal Analysis: S.P., J.-J.B., C.M.; Supervision: I.M., P.T.; Writing of Original Draft: A.P., P.T.; Review of Draft: all authors

## EXTENDED FIGURES

**Extended Figure 1.**
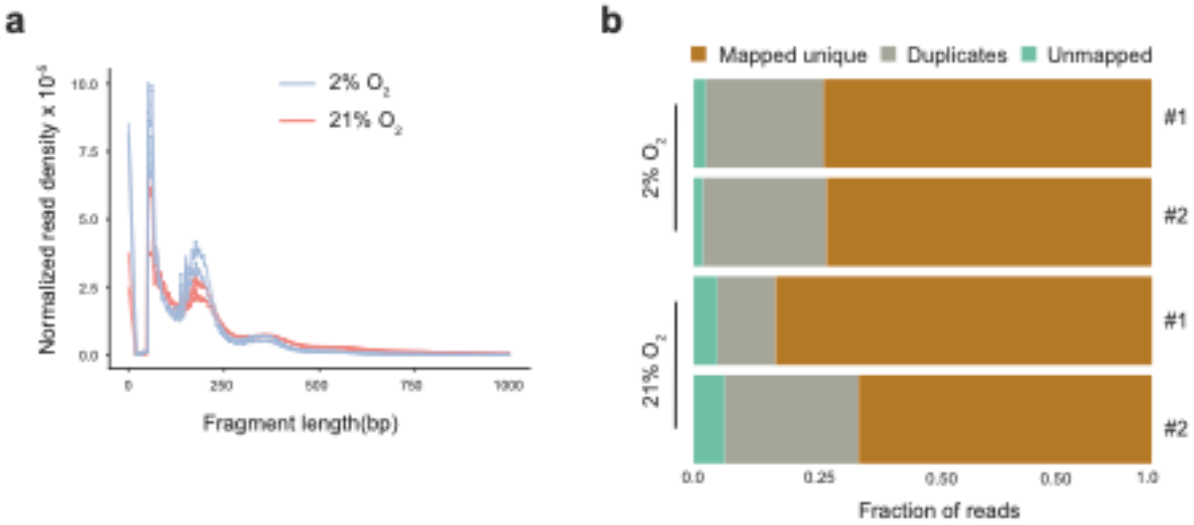
Quality control measurements of ATAC-seq. a) Insert size distribution of each single ATAC-seq library (2 libraries per oxygen condition-*a*) and b) mapping statistics for each individual replicate.

**Extended Figure 2.**
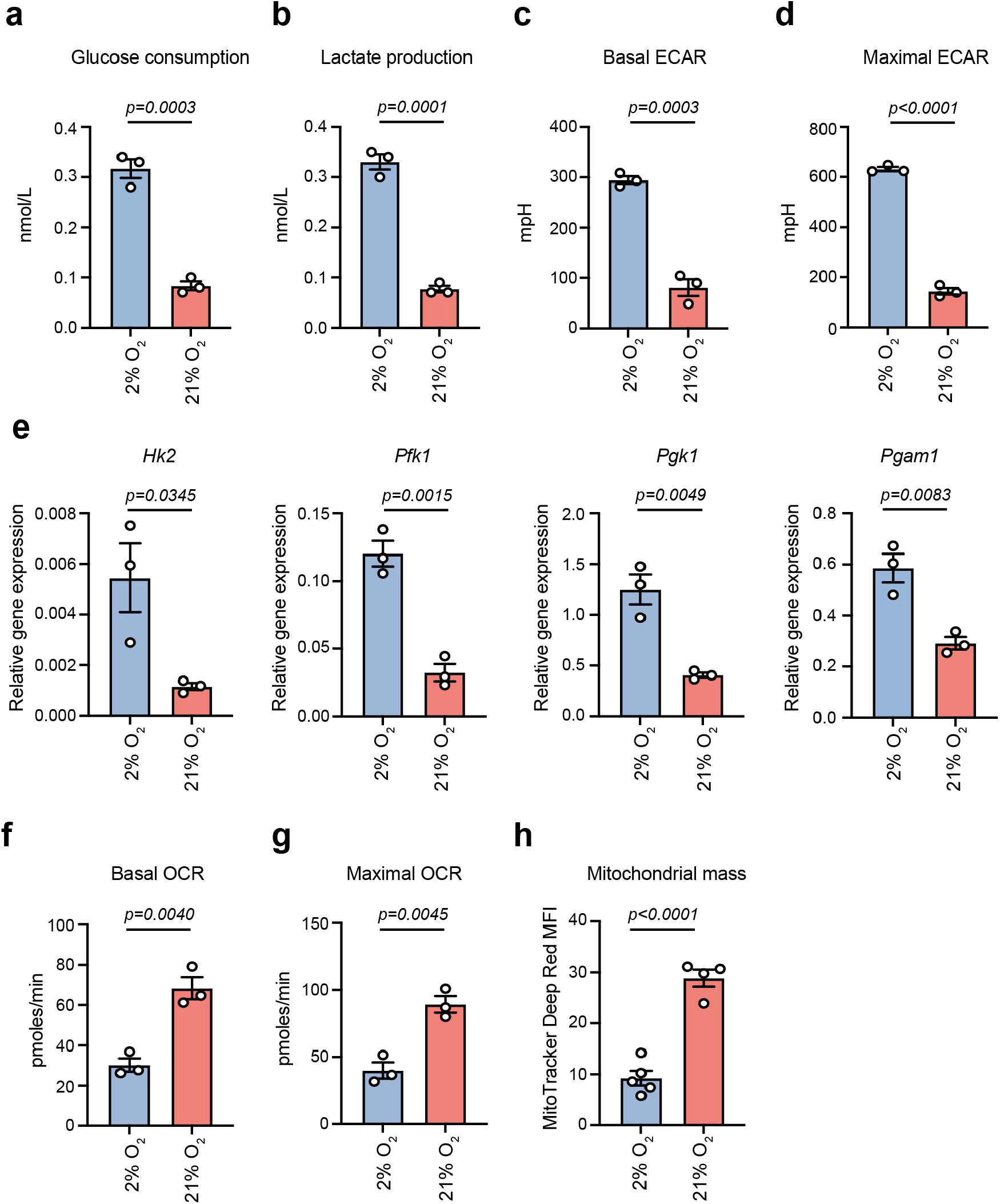
Metabolic characterization of normoxia-cultured cells. (**a-b**) Glucose consμmption (*a*) and lactate production (*b*) measured in the media of hypoxia- and normoxia-cultured cells using the Vi-Cell MetaFLEX instrμment. n=3 biologically independent experiments. (**c-d**) Basal (*c*) and maximal (*d*) ECAR in hypoxia- and normoxia-cultured BM-MSCs. n=3 biologically independent experiments. (**e**) qRT-PCR analysis of glycolytic genes. *β-actin* was used as an internal control for normalization. n=3 biologically independent experiments. (**f-g**) Basal (*f*) and maximal (*g*) OCR in hypoxia- and normoxia-cultured BM-MSCs. n=3 biologically independent experiments. (**h**) MFI of hypoxia- and normoxia-cultured cells after staining with the MitoTracker Deep Red FM dye. n = 4 biologically independent experiments.

**Extended Figure 3.**
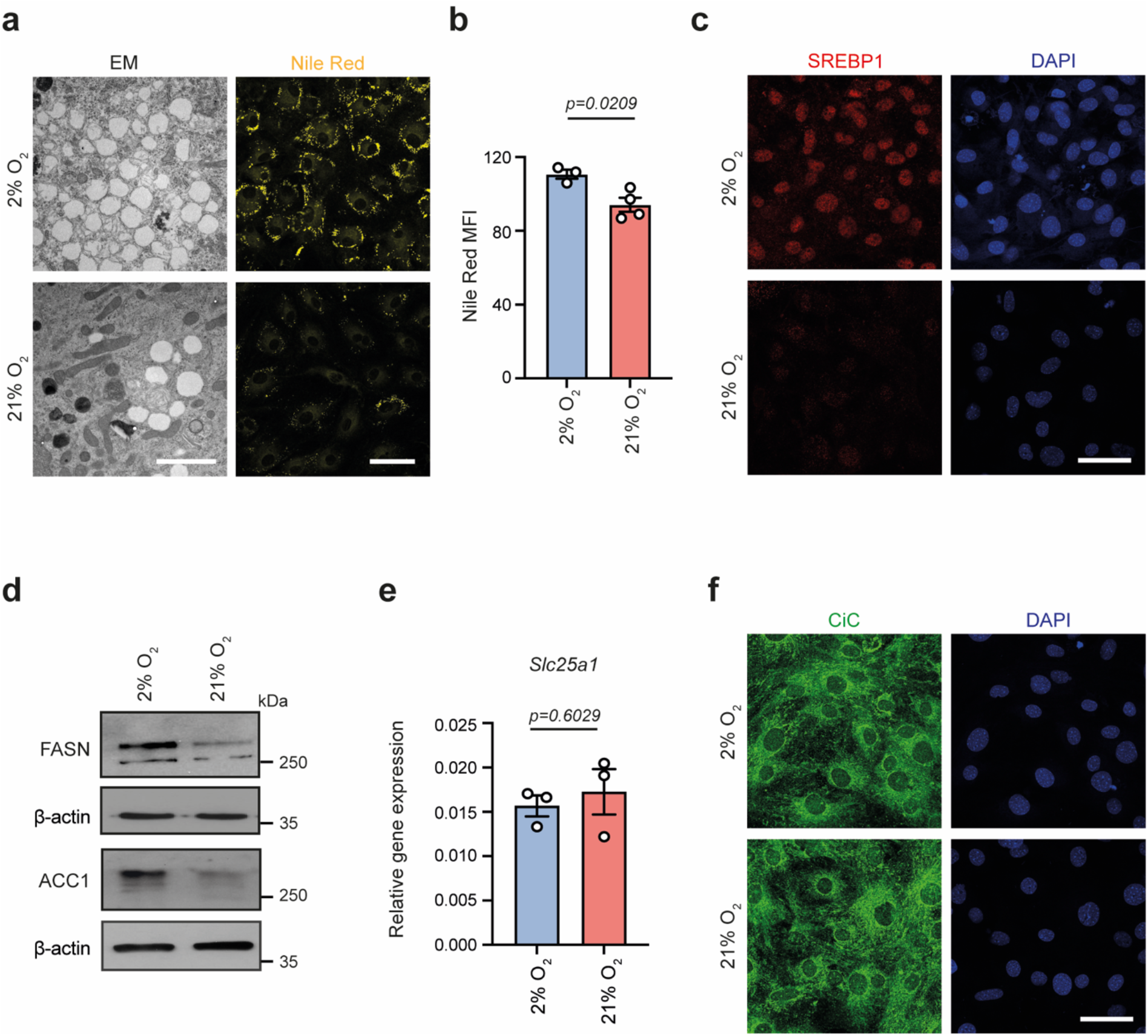
Impaired lipogenesis but stable CiC levels in normoxia-cultured cells. (**a-b**) Representative images (*a*) and quantification of lipid droplets after observing cells under the electron microscope (left) and after staining lipids with Nile Red (right-*b*). Scale bars, 2μm for electron microscopy images and 50μm for confocal images. (**c**) Representative images of hypoxia- and normoxia-cultured cells after immunostaining against SREBP1. Scale bar, 50μm. (**d**) Representative immunoblots for FASN and ACC1 in hypoxia- and normoxia-cultured cells. β-actin was used as loading control. n=3 biologically independent experiments. (**e**) qRT-PCR analysis of *Slc25a1*, which encodes citrate carrier. *β-actin* was used as an internal control for normalization. n=3 biologically independent experiments. (**f**) Representative images of hypoxia- and normoxia-cultured cells after immunostaining against CiC. Scale bar, 50μm.

